# CRISPR RNA-Guided Integrase Enables Targeted Genome Engineering and Prophage Functional Genomics in Plant-Pathogenic *Pseudomonas*

**DOI:** 10.64898/2026.09.22.753576

**Authors:** Cuong V. Hoang, Jackelene Cruz, Olakunle I. Olawole

## Abstract

*Pseudomonas* species include important plant pathogens responsible for substantial agricultural losses. Although CRISPR-INTEGRATE has been applied for genome engineering in medically relevant *Pseudomonas*, its utility in plant-pathogenic *Pseudomonas* has not been established. Here, we demonstrate CRISPR-INTEGRATE, an RNA-guided Tn7-like transposition system that enables targeted chromosomal integration without double-strand breaks or host-dependent homologous recombination, across three plant-pathogenic *Pseudomonas* backgrounds. Targeted disruption of *hrpJ*, *hrcC*, and *gacA* was confirmed by PCR and Sanger sequencing and resulted in distinct phenotypes, including impaired type III secretion-associated virulence and hypersensitive response, altered colony morphology and biofilm formation, and reduced motility. We further coupled CRISPR-INTEGRATE with Cre-*lox* recombination to excise a ∼40-kb prophage-associated region from the *P. syringae* pv. *tomato* DC3000 chromosome. Genomic characterization identified a 33.45-kb predicted prophage within this region, putative attL/attR sites sharing an identical 14-bp core sequence, an orthologous empty locus containing a candidate attB site, and conserved prophage-associated gene organization across *P. syringae* genomes. Deletion of the region altered bacterial growth and increased susceptibility to phage infection, with the strongest effect observed for phage Plaza, while having no detectable effect on the tested plant virulence phenotypes. Together, these findings establish CRISPR-INTEGRATE as a portable genome-engineering platform for plant-pathogenic *Pseudomonas* and demonstrate its utility for functional interrogation of individual genes and large accessory genomic elements.

## INTRODUCTION

*Pseudomonas syringae* is a Gram-negative plant pathogen comprising more than 60 pathovars collectively capable of infecting hundreds of plant species across major crop families, making it an economically important phytopathogen worldwide [1, 2]. Among characterized strains, *P. syringae* pv. *tomato* DC3000 (*Pst* DC3000) is a widely used model for studying plant–pathogen interactions. It infects both *Arabidopsis thaliana* and tomato (*Solanum lycopersicum*), has a fully sequenced genome, and possesses an extensively characterized repertoire of virulence effectors [2, 3]. *P. syringae* pv. *syringae* (Pss) has a similarly broad host range and is associated with blossom blight, shoot dieback, and frost injury in several agriculturally important hosts [2]. Beyond the *P. syringae* complex, *Pseudomonas viridiflava* is a widespread opportunistic plant pathogen associated with disease in diverse wild and cultivated plants [4, 5]. Together, these bacteria provide useful genetic backgrounds for investigating virulence, regulatory pathways, and accessory genome function in plant-pathogenic *Pseudomonas*.

Genetic manipulation is essential for defining gene function in *Pseudomonas*, but commonly used approaches have practical limitations. Allelic exchange using mobilizable suicide vectors typically requires cloning of flanking homology regions, conjugation, multiple rounds of selection, and *sacB*-based counterselection to recover defined chromosomal mutations [6]. The workflow can be labor-intensive, particularly for large genomic modifications. Mini-Tn7 systems provide stable, single-copy chromosomal integration and are widely used for complementation and reporter insertion, but integration is generally restricted to defined *attTn7* sites [7]. Recombineering approaches using lambda Red or RecTE-family recombinases can also be used for bacterial genome engineering but often require expression of heterologous recombination factors and optimization for different *Pseudomonas* backgrounds [8]. More recently, CRISPR-Cas9 and CRISPR-Cas12 systems have been adapted for *Pseudomonas*, frequently using double-strand break (DSB)-mediated counterselection to enrich for recombination-mediated genome edits [9, 10]. Although these approaches have expanded the genetic toolkit available for *Pseudomonas*, complementary methods that enable programmable chromosomal manipulation without DSB induction or dependence on host homologous recombination could further facilitate functional genomic studies.

CRISPR-INTEGRATE provides such an approach by coupling CRISPR RNA-guided target recognition with Tn7-like transposase-mediated DNA integration [11, 12]. The system directs insertion of user-defined DNA cargo at guide RNA-specified chromosomal sites without requiring DSB induction or host homology-directed repair. This mechanism enables programmable chromosomal targeting and provides a framework for multiplexed integration and manipulation of larger genomic regions. CRISPR-associated transposase systems have been demonstrated in several Gram-negative bacteria and applied to genome engineering in *Agrobacterium tumefaciens* [13, 14]. However, their utility for genome engineering in plant-pathogenic *Pseudomonas* has not been established.

Here, we evaluated CRISPR-INTEGRATE across three plant-pathogenic *Pseudomonas* backgrounds and targets representing different biological functions and genomic scales. We targeted the type III secretion system (T3SS), a major virulence determinant of phytopathogenic *Pseudomonas* encoded by the *hrp/hrc* gene cluster [15, 16]. Specifically, we disrupted *hrpJ* in *Pst* DC3000 and *hrcC* in Pss, providing well-characterized virulence-associated phenotypes for functional validation of genome editing [16, 17]. We further targeted *gacA* in *P. viridiflava*, a response regulator of the GacS/GacA two-component system that controls multiple virulence-associated traits in *Pseudomonas* [18, 19], to evaluate portability across species backgrounds. Finally, we coupled CRISPR-INTEGRATE with Cre-*lox* recombination to excise a ∼40-kb prophage-associated region from the *Pst* DC3000 chromosome [3]. Prophage-associated regions can contribute to bacterial fitness, phage interactions, and other accessory functions [20, 21], making their targeted manipulation useful for functional analysis of bacterial accessory genomes. Together, these experiments establish CRISPR-INTEGRATE as a programmable genome-engineering platform for plant-pathogenic *Pseudomonas* and demonstrate its application to both individual gene disruption and large-scale chromosomal deletion.

## RESULTS

### CRISPR-INTEGRATE architecture and validation of single-gene deletions across three Pseudomonas backgrounds

CRISPR-INTEGRATE couples RNA-guided target recognition by a Cascade-like ribonucleoprotein complex with transposase-mediated insertion of a donor mini-Tn cargo at a defined distance downstream of the guide RNA-specified protospacer (Fig. 1A). The complete editing system, comprising the TnsA–D transposition machinery, QCascade targeting complex, a BsaI-cloning CRISPR (CR) array, and mini-Tn donor flanked by right (R) and left (L) transposon ends, is encoded on the single delivery plasmid pKL2310 (Fig. 1B). We applied this platform to three chromosomal genes across three plant-pathogenic *Pseudomonas* backgrounds: *hrpJ* in *P. syringae* pv. *tomato* DC3000 (*Pst* DC3000), *hrcC* in *P. syringae* pv. *syringae*, and *gacA* in *P. viridiflava*. We subsequently extended the system to deletion of a ∼40-kb prophage-associated region in *Pst* DC3000. Delivery vectors carrying locus-specific guide RNAs were introduced into each strain by electroporation, and candidate mutants were screened by colony PCR using locus- and integration-specific primers (Fig. 2A). PCR analysis confirmed targeted cargo integration at *gacA*, *hrpJ*, and *hrcC* (Fig. 2B–D), and Sanger sequencing verified the expected integration junction (Fig. S1). These results demonstrate that the same CRISPR-INTEGRATE platform can mediate targeted chromosomal editing across three distinct plant-pathogenic *Pseudomonas* backgrounds.

**Figure 1.**
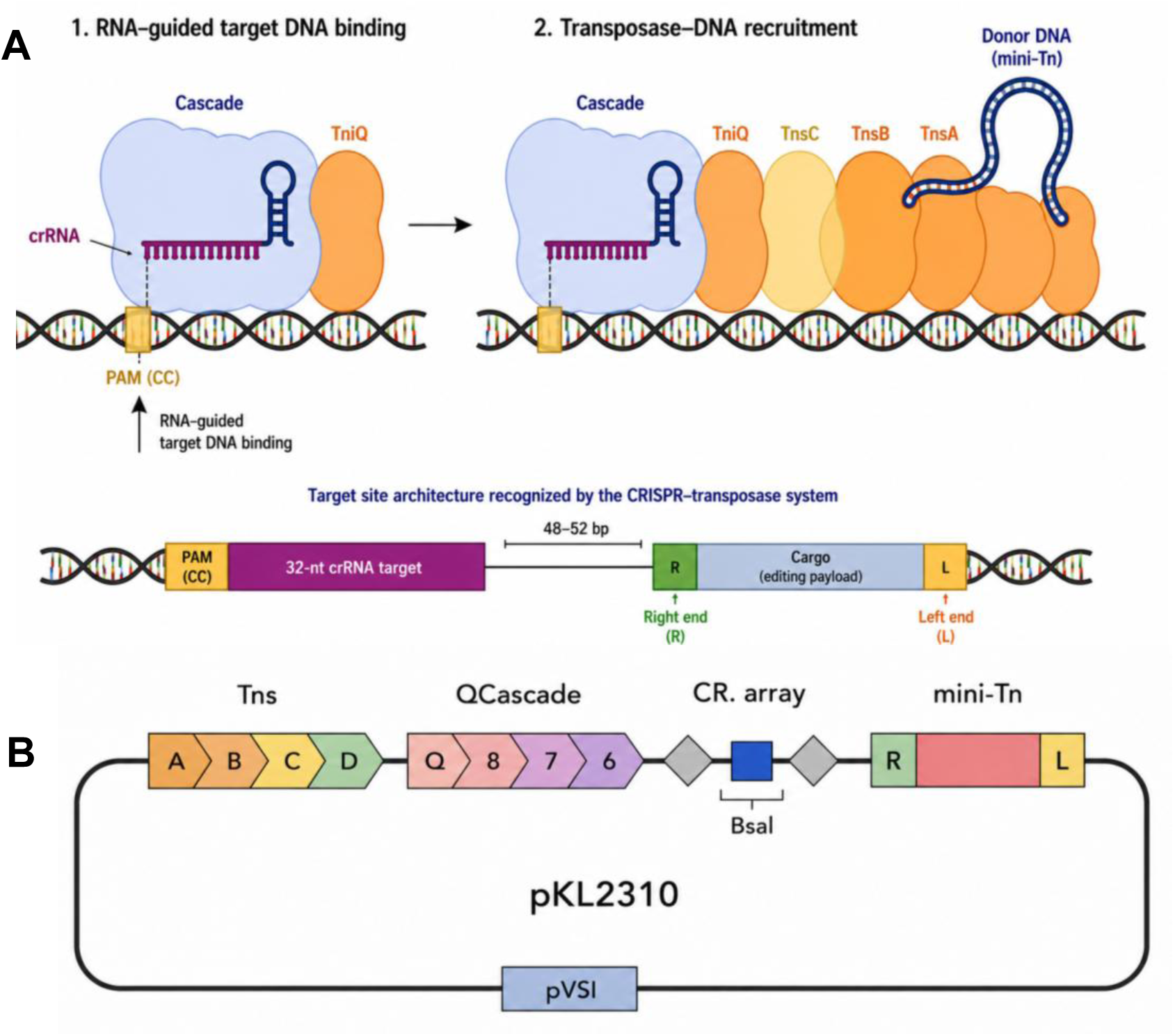
CRISPR-INTEGRATE platform architecture and experimental design. **(A)** Mechanism of RNA-guided transposition. The Cascade ribonucleoprotein complex, loaded with a crRNA and associated with TniQ, recognizes a target site by RNA-guided base-pairing adjacent to a CC PAM (left). Target recognition recruits the transposase complex (TniQ-TnsC-TnsB-TnsA), which mediates insertion of the donor mini-Tn cargo (right) at a fixed distance (48–52 bp) downstream of the 32-nt crRNA target site, flanked by defined transposon right (R) and left (L) ends (bottom). **(B)** Map of the CRISPR-INTEGRATE delivery plasmid pKL2310, encoding the Tns transposition genes (A–D), the QCascade targeting complex (Q, 8, 7, 6), a BsaI-cloning CRISPR (CR) array for spacer insertion, and the mini-Tn donor cargo (R–loxP–L) on a pVSI-derived broad-host-range backbone.

**Figure 2.**
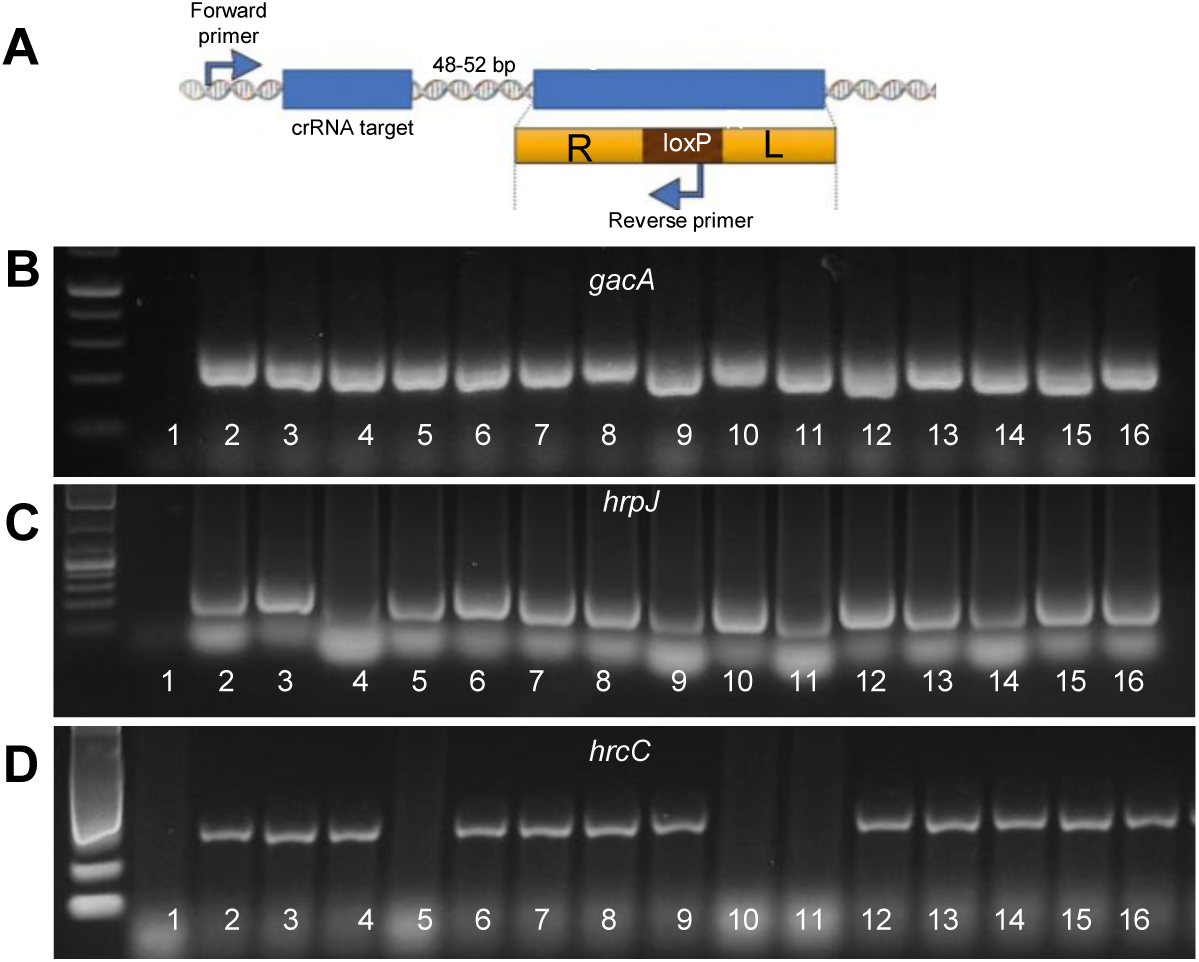
Genotypic confirmation of single-gene deletions. **(A)** Schematic of the colony PCR/Sanger sequencing strategy used to screen candidate mutants, showing the forward primer upstream of the crRNA target site, the 48–52 bp spacer to the integration site, and the reverse primer within the loxP-containing cargo. Representative colony PCR screen confirming precise CRISPR-INTEGRATE-directed deletion at *gacA* in *P. viridiflava* **(B)**, *hrpJ* in DC3000 **(C)**; *hrcC* in *P. syringae* pv. *syringae* **(D)**. Lanes 2-16 represent independent candidate colonies; lane 1 in each panel is the corresponding wildtype (*Pv*, DC3000, and *Pss*, respectively).

### Targeted disruption of *gacA*, *hrpJ*, and *hrcC* produces distinct regulatory and virulence phenotypes

Disruption of *gacA* in *P. viridiflava* resulted in loss of the mucoid colony morphology observed in the wild-type strain (Fig. 3A) and significantly reduced biofilm formation (Fig. 3B), consistent with the established role of GacA in regulating extracellular and surface-associated phenotypes in *Pseudomonas* [18, 19]. Infiltration assays further showed reduced lesion development in tomato and almond leaves inoculated with the *gacA* mutant compared with wild type at 3 days post-inoculation (dpi) (Fig. 3C–D). Disruption of the T3SS-associated genes *hrpJ* and *hrcC* similarly produced clear virulence-related phenotypes. Wild-type *Pst* DC3000 caused extensive disease symptoms in infiltrated tomato leaflets by 3 dpi, whereas the *hrpJ* mutant produced substantially reduced symptoms (Fig. 3E). Wild-type DC3000 also elicited a visible hypersensitive response (HR) in tobacco, whereas the *hrpJ* mutant produced no detectable cell-death response under the tested conditions (Fig. 3F). Likewise, wild-type *P. syringae* pv. *syringae* elicited necrosis on almond leaves and a clear HR in tobacco, whereas the *hrcC* mutant produced no visible response (Fig. 3G–H). These phenotypes are consistent with impaired T3SS function following disruption of *hrpJ* and *hrcC*. We next quantified bacterial motility to determine whether the targeted mutations affected an additional cellular phenotype. Motility was significantly reduced in the *gacA* (*P* = 0.0043), *hrcC* (*P* = 0.0053), and *hrpJ* (*P* = 0.0126) mutants relative to their respective parental strains (Fig. 3I). In contrast, deletion of the prophage-associated region described below did not significantly affect motility (*P* = 0.1912). Together, these phenotypic analyses provide functional validation of CRISPR-INTEGRATE-mediated editing at each single-gene target.

**Figure 3.**
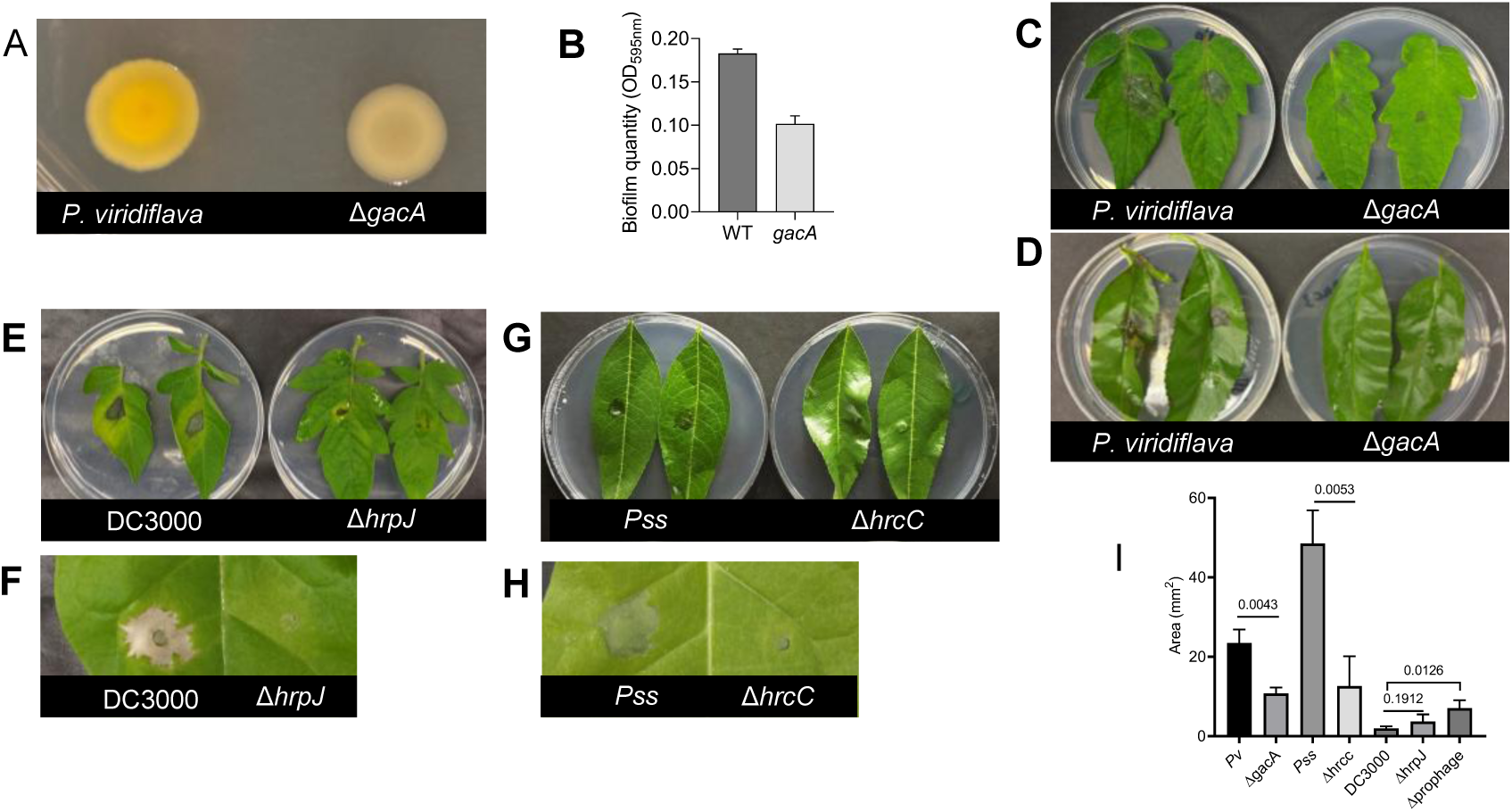
Phenotypic validation of CRISPR-INTEGRATE-generated deletion mutants. **(A)** Colony morphology of wild-type *P. viridiflava* (left) and the Δ*gacA* mutant (right) on LB agar. **(B)** Quantification of biofilm formation (OD₅₉₅) for wild-type and Δ*gacA* strains. **(C–D)** Disease symptoms in tomato and almond leaves infiltrated with wild-type *P. viridiflava* versus Δ*gacA* at 3 dpi. **(E)** Tomato leaves infiltrated with wild-type DC3000 and the Δ*hrpJ* mutant at 3 dpi. **(F)** Tobacco leaves infiltrated with wild-type DC3000 and Δ*hrpJ*, showing the hypersensitive response (HR) at 3 dpi. **(G)** Tomato leaf virulence assay comparing wild-type *P. syringae* pv. *syringae* and Δ*hrcC*. **(H)** Tobacco HR assay comparing wild-type *P. syringae* pv. *syringae* and Δ*hrcC* at 3 dpi. **(I)** Quantification of bacterial motility for each mutant relative to its parental wild-type strain across all four genetic backgrounds, with p-values indicated above each comparison.

### The DC3000 40-kb region is a conserved prophage-associated element with defined attachment-site architecture

We next characterized the ∼40-kb DC3000 region selected for large-scale genome engineering. Analysis of the experimentally targeted 40-kb interval identified a 33.45-kb prophage predicted by Phigaro within the broader region (Fig. 5A). Functional annotation revealed a structured phage-associated gene repertoire that included integration and excision functions, DNA metabolism, lysis and transcriptional regulation, together with tail, head, packaging, and connector proteins. Identical 14-bp sequences (TGGAAATCTTCAAA) were identified at or immediately adjacent to the experimental deletion boundaries (Fig. 4A). The left repeat was positioned adjacent to the integrase, whereas the right repeat occurred near the distal boundary of the element, supporting their designation as putative attL and attR sites. Comparison with the orthologous locus in *P. syringae* SUPP1331 provided additional evidence for this attachment-site architecture (Fig. 4B). SUPP1331 lacked the intervening ∼40.7-kb region and instead contained a single copy of the same 14-bp sequence at the corresponding chromosomal junction. The corresponding 100-bp host-facing flanks were 100% identical between DC3000 and SUPP1331, supporting the SUPP1331 junction as an orthologous empty locus containing a candidate attB site. Comparative synteny analysis further identified related prophage-associated regions in selected *P. syringae* genomes (Fig. 4C). TMP-centered regions from DC3000, UB303, SZ0049, 19B, Pss9097, and B48 showed substantial conservation of gene content and organization across the central region, with greater variability toward the flanks. Collectively, these analyses support the 40-kb DC3000 interval as a structurally organized prophage-associated element with identifiable attachment-site architecture and conserved, but variable, organization among *P. syringae* genomes.

**Figure 4.**
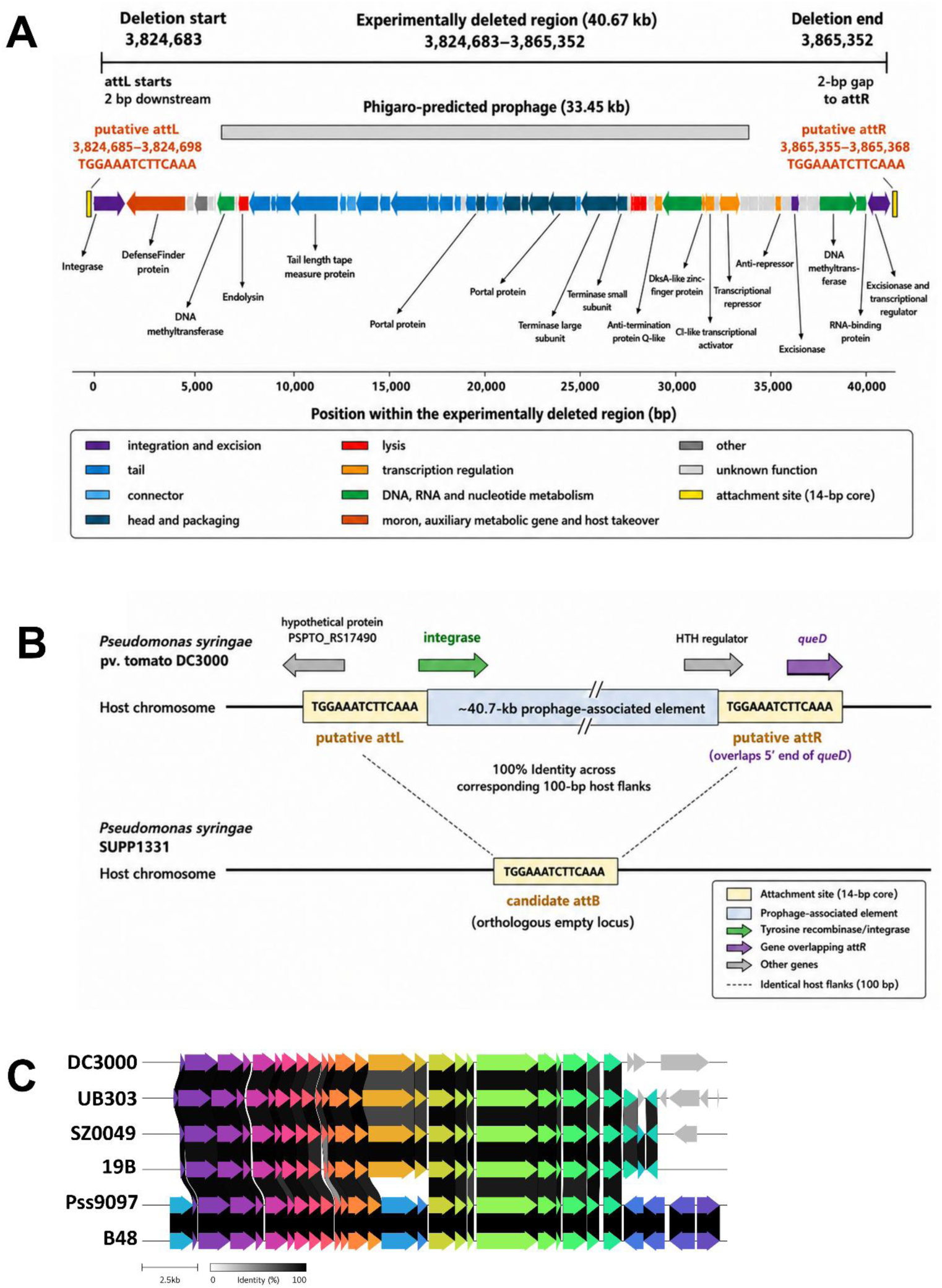
Genomic organization, attachment-site architecture, and comparative conservation of the DC3000 prophage-associated region. **(A)** Organization of the experimentally deleted 40-kb region in *Pseudomonas syringae* pv. *tomato* DC3000, showing the 33.45-kb Phigaro-predicted prophage, representative genes, functional categories, and putative attL/attR sites sharing an identical 14-bp core sequence. **(B)** Comparison with the orthologous empty locus in *P. syringae* SUPP1331, showing a single candidate attB site and 100% identity across the corresponding 100-bp host flanks. **(C)** Comparative synteny of tail tape-measure protein-centered prophage-associated regions from DC3000, UB303, SZ0049, 19B, Pss9097, and B48. Loci were oriented consistently and standardized around the tail tape-measure protein to facilitate comparison of gene organization. Homologous proteins are connected by shaded regions, with shading indicating amino acid sequence identity.

**Figure 5.**
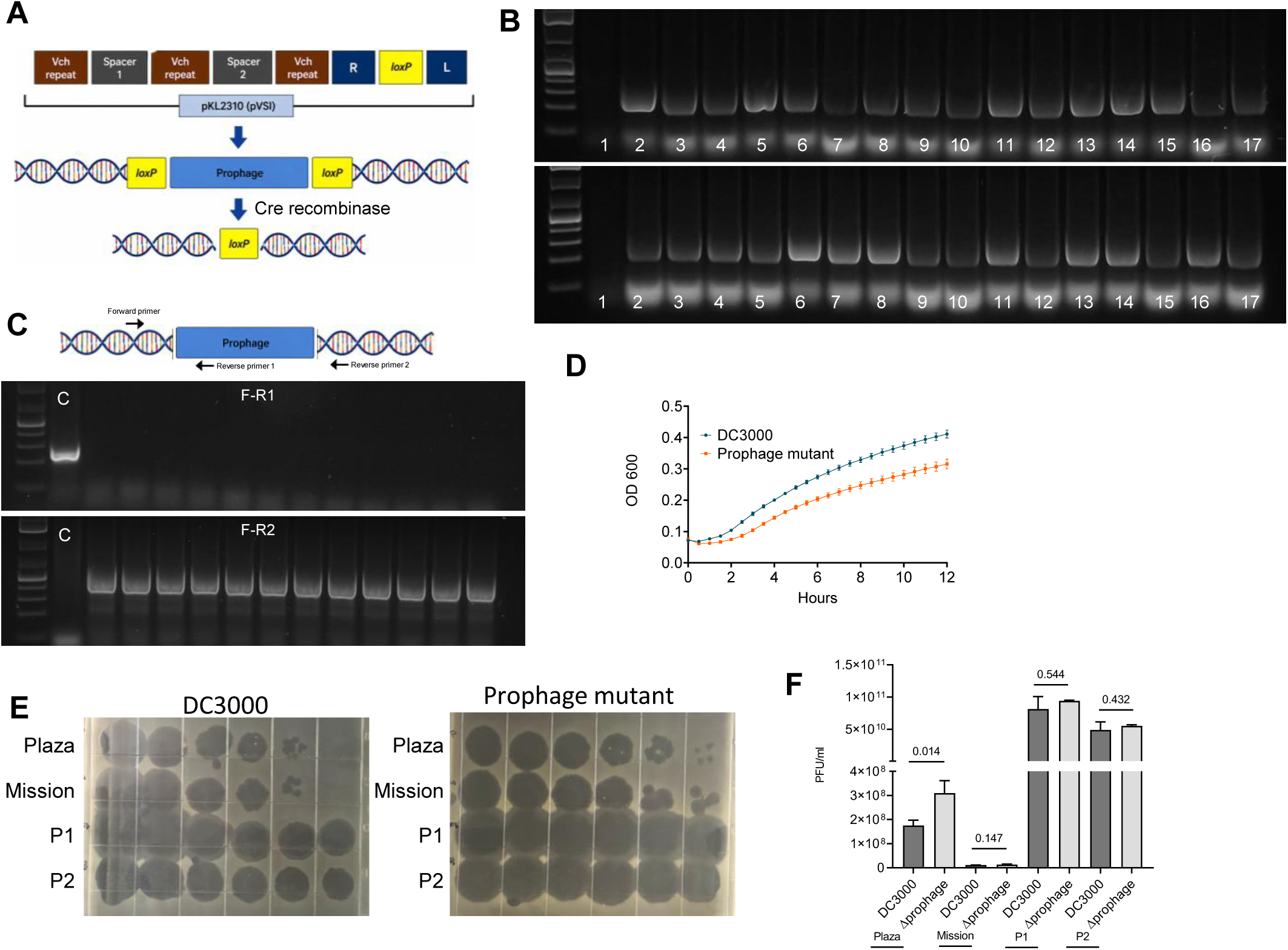
Two-step CRISPR-INTEGRATE/Cre-lox excision of a ∼40 kb prophage island in *Pst* DC3000. **(A)** Strategy overview: the deletion-cassette architecture on plasmid pKL2310, featuring two Vch-repeat/spacer units (Spacer 1 and Spacer 2) flanking a shared R–loxP–L cargo; CRISPR-INTEGRATE-directed integration of loxP-containing cassettes immediately upstream and downstream of the prophage island; and Cre recombinase-mediated excision of the intervening prophage sequence by recombination between the two loxP sites, leaving a single residual loxP scar. **(B)** Colony PCR screens confirming loxP cassette insertion at the upstream (top) and downstream (bottom) flanking integration sites. Lanes 2-17 represent independent candidate colonies; lane 1 in each panel is the corresponding wildtype (DC3000). **(C)** Colony PCR screens following Cre-mediated excision, C represents bacteria carrying two loxP sites. **(D)** Growth curve (OD₆₀₀ over 12 h) comparing wild-type *Pst* DC3000 and the prophage deletion mutant in liquid culture. **(E)** Phage spot-titer assay comparing phage susceptibility of wild-type DC3000 (top) and the prophage deletion mutant (bottom) across a dilution series; rows 1–4 represent phage Plaza, Mission, P1, and P2, showing increased plaque formation in the deletion mutant. **(F)** Quantification of phage titer (PFU/ml) for wild-type DC3000 and the prophage deletion mutant against four phage isolates (Plaza, Mission, P1, and P2), with p-values indicated above each comparison.

### CRISPR-INTEGRATE/Cre-lox enables excision of the 40-kb prophage-associated region in *Pst* DC3000

We next tested whether CRISPR-INTEGRATE could be extended from single-gene editing to large-scale chromosomal engineering by deleting the 40-kb prophage-associated region from *Pst* DC3000. Deletion was achieved using a two-step CRISPR-INTEGRATE/Cre-lox strategy. Two guide RNAs directed insertion of loxP-containing mini-Tn cassettes at sites flanking the targeted region, followed by Cre-mediated recombination between the two loxP sites to excise the intervening sequence (Fig. 5A). Following electroporation and selection, colony PCR identified clones carrying the expected upstream and downstream loxP integrations (Fig. 5B). Following Cre expression, candidate deletion mutants produced the expected excision-junction amplicon (Fig. 5C). In contrast, PCR targeting an internal sequence within the 40-kb region produced an amplicon from wild-type DNA but not from the deletion mutant, further confirming loss of the targeted interval. Sanger sequencing of the deletion junction verified joining of the flanking chromosomal sequences at the designed loxP recombination site (Fig. S2). These results demonstrate that CRISPR-INTEGRATE can be coupled with Cre-lox recombination to engineer large chromosomal deletions in *Pst* DC3000.

### Deletion of the prophage-associated region alters bacterial growth and phage susceptibility

We next examined the biological consequences of deleting the prophage-associated region. The deletion mutant displayed altered growth relative to wild-type *Pst* DC3000, reaching a lower final OD₆₀₀ during the 12-h growth assay (Fig. 5D). In contrast, motility was not significantly different between the deletion mutant and wild type (*P* = 0.1912; Fig. 3I). We previously characterized the lytic phages Mission, Mobley, and Plaza, which infect almond-associated *Pseudomonas* strains [22]. To determine whether deletion of the prophage-associated region altered susceptibility to phage infection, wild-type DC3000 and the deletion mutant were challenged with these phages and other selected phages. Spot-titer assays showed greater plaque formation on the deletion mutant than on wild-type DC3000 across the tested phages (Fig. 5E). Quantification showed a significant difference for Plaza (*P* = 0.014), whereas Mission (*P* = 0.147), P1 (*P* = 0.544), and P2 (*P* = 0.432) showed similar directional trends that did not reach statistical significance (Fig. 5F). Thus, deletion of the 40-kb region increased susceptibility most clearly to Plaza, while evidence for effects on the other phages was more limited. Despite these growth and phage-susceptibility phenotypes, the deletion mutant remained comparable to wild-type DC3000 in the tested plant virulence and HR assays (Fig. S3), indicating that the region was not required for these phenotypes under the experimental conditions examined. Together, these findings demonstrate the utility of CRISPR-INTEGRATE for engineering large accessory-genome regions and suggest that the DC3000 prophage-associated element contributes to specific bacterial fitness and phage-interaction phenotypes.

## DISCUSSION

### CRISPR-INTEGRATE expands functional genome engineering in plant-pathogenic *Pseudomonas*

This study extends CRISPR RNA-guided transposition to plant-pathogenic *Pseudomonas* and demonstrates that the same core system can function across distinct phytopathogenic backgrounds. CRISPR-associated transposases differ fundamentally from nuclease-based CRISPR systems because RNA-guided DNA recognition is coupled to transposition rather than double-strand DNA cleavage [11, 12]. Consequently, targeted integration does not require repair of a programmed double-strand break or homologous recombination at the target locus. The original INTEGRATE platform demonstrated efficient and multiplexed genome engineering in several Gram-negative bacteria, including *Pseudomonas putida* [23]. Our findings extend this capability to plant-pathogenic *Pseudomonas* and show that a common editing framework can be applied across bacterial backgrounds differing in host association, regulatory circuitry, and genomic composition. The biological significance of this portability is illustrated by disruption of *hrpJ*, *hrcC*, and *gacA*. The T3SS is central to *P. syringae*–plant interactions, and the phenotypes associated with *hrpJ* and *hrcC* disruption were consistent with their established roles in T3SS-dependent pathogenicity and host responses [24, 25]. Thus, the value of these mutants extends beyond confirmation of targeted integration: they demonstrate that RNA-guided transposition can generate functionally informative perturbations of major virulence pathways in planta. The accompanying changes in motility suggest additional phenotypic consequences of these disruptions, although whether these effects are direct or secondary remains unresolved.

The *gacA* mutant further demonstrates the utility of this approach for interrogating global regulatory networks. GacS/GacA regulates diverse traits in *Pseudomonas*, including motility, biofilm formation, secondary metabolism, stress responses, and virulence [26]. The combined changes in colony morphology, biofilm formation, motility, and plant-associated phenotypes following *gacA* disruption in *P. viridiflava* are consistent with perturbation of a regulatory system coordinating multiple bacterial lifestyles. Similar GacA-dependent phenotypes have been reported in other *Pseudomonas* backgrounds [27], although the regulatory consequences can vary substantially among strains and species [28]. This context dependence highlights a broader application of CRISPR-INTEGRATE: systematic perturbation of conserved genes across diverse *Pseudomonas* backgrounds could help distinguish core regulatory functions from lineage- or niche-specific adaptations. The present work is conceptually related to the application of CRISPR-INTEGRATE in *Agrobacterium tumefaciens*, where RNA-guided integration was used for single and multiplex gene disruption and, together with Cre-*lox*, for large chromosomal deletions [13, 14]. The significance of our study therefore lies not in establishing these capabilities for the first time, but in extending them to plant-pathogenic *Pseudomonas* and applying them to biologically distinct targets ranging from virulence and regulatory genes to a large accessory genomic element.

### Genomic architecture supports the DC3000 40-kb region as a dynamic prophage-associated element

Characterization of the large deletion target revealed that the experimentally defined ∼40-kb region has multiple features consistent with a biologically integrated prophage-associated element. Although Phigaro predicted a smaller 33.45-kb prophage within the interval, the broader region contains recognizable structural, packaging, lysis, regulatory, integration, and excision functions. The presence of an integrase near one boundary and identical 14-bp repeats at or immediately adjacent to the two boundaries further supports this interpretation. The orthologous locus in SUPP1331 provides additional evidence for the proposed attachment-site architecture. Whereas DC3000 contains two copies of the 14-bp sequence flanking the prophage-associated interval, SUPP1331 lacks the intervening element and contains a single copy at the corresponding chromosomal junction. Conservation of the adjacent host sequences is consistent with these repeats representing putative attL and attR sites in DC3000 and the SUPP1331 junction representing a candidate empty attB site. Nevertheless, demonstrating excision, circularization, or productive induction will be necessary to establish whether the DC3000 element remains an active inducible prophage rather than an integrated prophage remnant.

Comparative synteny further suggests that this region is an evolutionarily dynamic component of the *P. syringae* accessory genome. Conservation of a central phage structural module among selected strains, together with greater variation toward the flanks, is consistent with the modular evolution characteristic of phage-associated regions. Prophage diversity has similarly been implicated in genome diversification and adaptation within *P. syringae* [29], while other mobile genomic islands in this species can be maintained or lost depending on selective conditions [30]. The genomic architecture of the DC3000 region therefore provided a strong rationale for moving beyond sequence-based prediction to direct experimental interrogation of its biological function.

### Prophage deletion links accessory-genome content to bacterial fitness and phage interactions

The altered growth and phage susceptibility of the deletion mutant indicate that the DC3000 prophage-associated region is not simply a passive component of the chromosome. Prophages can modify host physiology and provide protection against subsequent phage infection through mechanisms acting at adsorption, DNA entry, replication, or later stages of phage development [31]. The increased susceptibility of the deletion mutant, particularly to Plaza, is consistent with the possibility that one or more functions within the deleted region influence phage infection. Importantly, this effect was phage dependent. Plaza showed a statistically significant difference, whereas Mission, P1, and P2 showed directional but nonsignificant differences. The region should therefore not yet be considered a general superinfection-immunity locus. Instead, the phenotype suggests that it influences susceptibility to particular phages, potentially through mechanisms that differ according to phage biology. Comparing adsorption, efficiency of plating, intracellular replication, and burst size between the parental and deletion strains could identify the stage of infection affected and help distinguish receptor-mediated effects from intracellular defense.

The growth phenotype further suggests that the biological contribution of the region extends beyond phage interactions. Prophage carriage can impose fitness costs or provide benefits depending on bacterial genotype and environmental conditions [32]. The reduced growth of the deletion mutant is therefore consistent with a positive contribution of one or more genes within the region to bacterial physiology under the conditions examined. Because the deletion removed a large multigene interval, however, the phenotype cannot presently be assigned to a specific prophage function. Conversely, deletion did not produce a detectable effect on the tested plant virulence or HR phenotypes. This suggests that the contribution of the region is separable from the major T3SS-dependent pathogenicity functions measured here. Accessory elements need not produce pronounced changes in acute disease symptoms to be biologically important; their effects may instead become apparent during phage exposure, competition, environmental stress, epiphytic survival, or long-term host colonization. The combined growth and phage-susceptibility phenotypes therefore point toward a role in bacterial fitness and phage interactions rather than an essential contribution to the plant disease phenotypes examined.

### Large-region engineering connects comparative genomics with experimental accessory-genome biology

A broader implication of this study is that CRISPR-INTEGRATE can facilitate functional interrogation of bacterial genomes at multiple scales. Comparative genomics has revealed extensive accessory-genome diversity in plant-pathogenic *Pseudomonas*, but assigning biological functions to large genomic regions remains difficult when genetic analysis proceeds one gene at a time. Prophages, genomic islands, and other mobile elements may contain dozens of genes whose collective contribution cannot readily be inferred from sequence annotation alone. Coupling RNA-guided *loxP* placement with Cre-mediated excision provides a useful strategy for addressing this problem. An entire candidate region can first be removed to determine whether it contributes to a measurable phenotype, after which smaller deletions, individual gene disruptions, or complementation can localize the responsible functions. Viewed in this context, the ∼40-kb deletion is not primarily a demonstration of deletion size. Rather, it represents a functional-genomics experiment in which comparative genomic predictions were converted into an experimentally testable phenotype. The resulting phage-susceptibility phenotype now provides a foundation for genetically dissecting the contribution of individual genes or modules within the region.

This application also distinguishes the present study from previous implementation of INTEGRATE in *A. tumefaciens*, where large Cre-*lox*-mediated deletions, including a 42.6-kb interval, were already demonstrated [13, 14]. Here, large-region engineering was applied to a naturally occurring prophage-associated element and integrated with comparative genomics and phenotypic analysis. This combination provides a framework for experimentally testing accessory-genome functions that otherwise remain largely inferred from genomic sequence.

### Implications, limitations, and future applications

Several questions remain before the broader capabilities of CRISPR-INTEGRATE in phytopathogenic *Pseudomonas* can be defined. Although targeted integrations and the large deletion were confirmed by junction PCR and Sanger sequencing, whole-genome sequencing would provide a more comprehensive assessment of editing specificity and potential secondary genomic changes. Likewise, successful editing in three genetic backgrounds supports portability but does not establish universal applicability across *Pseudomonas*. Transformation efficiency, PAM availability, target accessibility, plasmid maintenance, and strain-specific interactions with the transposition machinery may influence editing outcomes. The prophage-deletion phenotype also creates a tractable path toward mechanistic analysis. Progressive sub-deletions and complementation of candidate genes could identify the genetic determinants underlying altered growth and phage susceptibility, while adsorption and intracellular phage replication assays could determine the affected stage of infection. Prophage induction and circularization experiments would further establish whether the element retains the capacity for active excision or productive development.

More broadly, RNA-guided integrases can support multiplexed bacterial genome engineering [13, 14], providing opportunities to perturb combinations of virulence genes, regulatory pathways, or accessory loci. Such capabilities could be particularly valuable in *Pseudomonas*, where functional redundancy and extensive accessory-genome variation can obscure phenotypes produced by individual mutations. Overall, our findings establish CRISPR-INTEGRATE as a portable genome-engineering approach for plant-pathogenic *Pseudomonas* and demonstrate its value as a tool for functional genomics rather than simply mutant construction. By enabling targeted perturbation of individual genes and experimental removal of large accessory regions, the platform provides a means to connect comparative genomic predictions with causal biological phenotypes. The DC3000 prophage-associated region illustrates this potential: genomic evidence supports its identity as an evolutionarily variable accessory element, while its deletion reveals contributions to bacterial physiology and phage susceptibility without a detectable requirement for the tested plant pathogenicity phenotypes. CRISPR-INTEGRATE therefore provides a framework for investigating how virulence determinants, regulatory networks, prophages, and other accessory genomic elements shape the biology and evolution of plant-pathogenic *Pseudomonas*.

## MATERIALS AND METHODS

### Bacterial strains, plasmids, and growth conditions

*Pseudomonas syringae* pv. *tomato* DC3000 (*Pst* DC3000), *P. syringae* pv. *syringae* (*Pss*), and *Pseudomonas viridiflava* were used as wild-type strains throughout this study. All *Pseudomonas* strains were routinely cultured on Luria-Bertani (LB) medium at 28°C with shaking at 200 rpm. *Escherichia coli* DH5α was used for plasmid propagation and maintained on LB medium at 37°C. Where required for plasmid maintenance or selection of chromosomal mutants, spectinomycin and kanamycin were added at 50 µg/ml. A Cre recombinase expression plasmid (pKL2315) was additionally used to catalyze loxP-mediated excision of the prophage island.

### CRISPR-INTEGRATE guide RNA design and vector construction

Guide RNAs (gRNAs) targeting each locus were designed using EuPaGDT (http://grna.ctegd.uga.edu/) to identify protospacer sequences adjacent to the CRISPR-INTEGRATE PAM sequence (CC). gRNAs were selected to minimize predicted off-target sites within the respective *Pseudomonas* genomes. For individual gene deletions, the gRNA spacer sequence was cloned into the CRISPR-INTEGRATE delivery vector pKL2310 by BsaI digestion and ligation. For the prophage-island deletion, two gRNAs targeting sequences immediately upstream (Spacer 1) and downstream (Spacer 2) of the prophage region were cloned into pKL2310. Successful spacer insertion into pKL2310 was verified by colony PCR using a gene-specific forward primer and 2310-loxP-R. All oligonucleotide sequences used for gRNA cloning, cassette construction, and screening PCR are listed in Supplementary Table S2.

### Genomic characterization and comparative analysis of the DC3000 prophage-associated region

The experimentally targeted ∼40-kb region of the *Pseudomonas syringae* pv. *tomato* DC3000 chromosome was characterized using complementary prophage prediction, functional annotation, attachment-site analysis, and comparative genomics. Prophage boundaries within the region were predicted using Phigaro [33], and coding sequences were functionally characterized using Pharokka [34] and PHOLD [35]. Predicted proteins were grouped into functional categories, including integration/excision, structural and packaging functions, lysis, transcriptional regulation, and DNA/RNA metabolism. The resulting annotations were used to generate a gene map of the experimentally targeted region.

To identify candidate prophage attachment sites, sequences surrounding the experimental deletion boundaries were examined for direct repeats. An identical 14-bp sequence (TGGAAATCTTCAAA) was identified at both boundaries and designated putative attL and attR based on its genomic position and proximity to the predicted integrase. The corresponding chromosomal region in *P. syringae* SUPP1331 was subsequently examined as a candidate empty locus. The DC3000 host-facing sequences flanking the prophage-associated region were aligned to the orthologous SUPP1331 region, and sequence identity across the corresponding 100-bp flanks was determined. The presence of a single copy of the 14-bp sequence at the SUPP1331 junction was used to define a candidate attB site.

For comparative analysis, the DC3000 prophage-associated sequence was searched against available *P. syringae* genome sequences using BLASTN to identify related loci. Representative loci spanning different levels of sequence conservation were selected from DC3000, UB303, SZ0049, 19B, Pss9097, and B48. Genomic regions were oriented consistently and standardized around the predicted tail tape-measure protein (TMP) to facilitate comparison. Gene content, order, and protein homology were compared using Clinker, with homologous proteins connected according to amino acid sequence similarity.

### Construction of chromosomal deletion mutants

CRISPR-INTEGRATE delivery vectors were introduced into each *Pseudomonas* strain by electroporation. Overnight cell culture was harvested by centrifugation, washed three times with ice-cold sterile water, and resuspended in 10% glycerol. Electroporation was performed using 100 µl of electrocompetent cells and 200 ng of plasmid DNA in a 0.2 cm cuvette at 2.5 kV, 25 µF, and 200 Ω. Cells were recovered in 500 µl of 2x LB medium at 28°C for 2 hours with shaking and plated on LB agar supplemented with antibiotics. Colonies were screened by colony PCR using gene-specific forward primer and 2310-loxP-R, and candidate mutants exhibiting the expected deletion-specific band were confirmed by Sanger sequencing.

### Deletion of the prophage-like genomic island in *Pst* DC3000

A ∼40-kb prophage-like genomic island on the *P. syringae* pv. tomato DC3000 chromosome [3] was excised using a two-step CRISPR-INTEGRATE/Cre-lox strategy rather than by direct single-step deletion. A CRISPR-INTEGRATE construct carrying Spacer 1 and Spacer 2 was introduced into *Pst* DC3000 by electroporation to insert loxP sites upstream and downstream of the prophage-like region (Fig. 5A). Following sequence confirmation of both flanking loxP insertions, Cre recombinase was supplied in trans using pKL2315 to catalyze site-specific recombination between the two loxP sites. This recombination excised the intervening ∼40-kb prophage-like sequence, leaving a single residual loxP scar at the recombination junction (Fig. 5A).

### Plant virulence assay

To assess virulence, wild-type and deletion-mutant strains were infiltrated into detached leaves of 4-week-old tomato (*Solanum lycopersicum*) and almond (*Prunus dulcis*) plants. Tomato and almond leaves were maintained on agar plates under a 16-h light/8-h dark photoperiod at 22°C. Bacterial strains were grown overnight in LB medium, harvested by centrifugation, washed twice with sterile 10 mM MgCl₂, and resuspended in 10 mM MgCl₂ to an OD₆₀₀ of 0.4. Bacterial suspensions were infiltrated into the abaxial surface of fully expanded leaves using a needleless 1-mL syringe. At least six leaves were inoculated per strain in each experiment. Disease symptoms were evaluated and photographed at 3 days post-inoculation (dpi). Experiments were performed at least twice with consistent results.

### Hypersensitive response assay

Wild-type and deletion mutant strains were tested for HR elicitation in *Nicotiana tabacum*. Bacterial strains were prepared as described above and resuspended to OD₆₀₀ of 0.2 in 10 mM MgCl₂. Suspensions were infiltrated into fully expanded tobacco leaves. HR development was assessed visually at 3 dpi. Experiments were repeated three times independently.

### Swimming mobility assay

Bacterial strains were grown overnight in 5 mL LB medium at 28°C with shaking at 200 rpm. Overnight cultures were standardized to an OD₆₀₀ of 0.4. Swimming assays were performed on LB medium containing 0.3% soft agar. Approximately 25 mL of molten medium was poured into sterile 90-mm Petri dishes and allowed to dry with the lids partially open for 1 h before inoculation. A sterile inoculation loop was used to inoculate each plate by stabbing approximately halfway into the center of the soft-agar surface. Plates were incubated upside down at 28°C. Images were captured at 24 h using a fixed overhead imaging setup, with a ruler included as a size reference. Swimming motility was quantified using ImageJ by calibrating images to millimeters, converting images to 8-bit grayscale, and adjusting the threshold to generate a binary image. The area of the swimming zone was measured in mm² using the Analyze Particles function. The threshold was manually adjusted when necessary to accurately define the boundary of the motility zone.

## Acknowledgments

This study was supported by the initial complement package provided to O.O. by the College of Natural and Agricultural Sciences, University of California, Riverside.

## Author contributions

O.I.O. conceived and supervised the study. C.V.H. and J.C. performed the experiments. C.V.H. prepared the original manuscript draft, and O.I.O. reviewed and edited the manuscript. All authors read and approved the final manuscript.

## Supplemental Figure Legends

**Table S1.** Efficiency of CRISPR-INTEGRATE-directed genome editing across target loci.

**Figure S1. Sanger sequencing validation of CRISPR-INTEGRATE-mediated integration at the *gacA* locus.** Sanger sequencing chromatogram confirming integration of the CRISPR-INTEGRATE cassette at the targeted *gacA* locus. The *gacA* crRNA target site, CC PAM, and the integration site located 48 bp downstream of the crRNA target site are indicated. The light-blue region represents the integrated R–loxP–L cassette. The target site duplication (TSD) and flanking *gacA* sequences are also indicated, confirming the expected integration at the *gacA* locus.

**Figure S2. Sanger sequencing validation of prophage deletion.** Sanger sequencing chromatograms confirming deletion of the targeted prophage region. The top trace shows the upstream prophage boundary, with the integration site 50 bp downstream of crRNA site 1; the bottom trace shows the downstream boundary, with the integration site 49 bp downstream of crRNA site 2. The prophage sequences were removed following Cre-mediated excision, leaving a single loxP site at the deletion junction. The sequencing traces show the regions flanking the deleted prophage and the remaining loxP sequence, confirming the expected prophage deletion.

**Figure S3. Phenotypic characterization of the prophage deletion mutant. (A)** Tomato leaves infiltrated with wild-type DC3000 and the prophage deletion mutant at 3 days post-infiltration (dpi). **(B)** Tobacco leaves infiltrated with wild-type DC3000 and the prophage deletion mutant, showing the hypersensitive response (HR) at 3 dpi. No differences in virulence or HR were observed between wild-type DC3000 and the prophage deletion mutant, confirming that the prophage-like island is dispensable for T3SS-mediated pathogenicity functions under the tested conditions.

**Table S2.** List of oligonucleotides used in this study

